# Selection for Intermediate Genotypes Enables a Key Innovation in Phage Lambda

**DOI:** 10.1101/018606

**Authors:** Alita R. Burmeister, Richard E. Lenski, Justin R. Meyer

## Abstract

The evolution of qualitatively new functions is fundamental for shaping the diversity of life. Such innovations are rare because they require multiple coordinated changes. We sought to understand the evolutionary processes involved in a particular key innovation, whereby phage λ evolved the ability to exploit a novel receptor, OmpF, on the surface of *Escherichia coli* cells. Previous work has shown that this transition repeatedly evolves in the laboratory, despite requiring four mutations in specific regions of a single gene. Here we examine how this innovation evolved by studying six intermediate genotypes that arose during independent transitions to use OmpF. In particular, we tested whether these genotypes were favored by selection, and how a coevolved change in the hosts influenced the fitness of the phage genotypes. To do so, we measured the fitness of the intermediate types relative to the ancestral λ when competing for either ancestral or coevolved host cells. All six intermediates had improved fitness on at least one host, and four had higher fitness on the coevolved host than on the ancestral host. These results show that the evolution of the phage’s new ability to use OmpF was repeatable because the intermediate genotypes were adaptive and, in many cases, because coevolution of the host favored their emergence.

## Introduction

A striking feature of the living world is the great variety of functions that organisms have evolved, but a detailed understanding of how species have acquired specific functions is difficult because of limitations of both theory and data (Gavrilets 2010). One challenge is to understand how populations evolve traits that require multiple interacting changes, such as specific deformations in the reactive pocket of an enzyme (Voordeckers et al. 2012) or the specialized modifications of an appendage (Moczek and Rose 2009). If each intermediate state leading to the new function is more fit than its immediate predecessor, then it can evolve readily, even if the fitness benefits do not accrue from the new function *per se* (Lenski et al. 2003). By contrast, the evolution of a new function will be slower and more difficult – although not impossible (Covert et al. 2013) – if any of the requisite intermediates have lower fitness than their progenitors.

Meyer *et al.* (2012) reported observing the evolution of a new function in action. While studying populations of bacteriophage λ propagated in the laboratory, the authors found that λ sometimes evolved to target a new receptor on the surface of its host, the bacterium *Escherichia coli*. Phage λ typically attaches to a porin protein called LamB; however, some evolved genotypes could also use a different porin called OmpF. This new function had not previously been observed, despite decades of intensive study of λ by virologists and molecular biologists. One reason this new function had not previously been observed was that it required four mutations in a gene, called *J*, that encodes a tail protein, J, that λ uses to recognize and attach to the receptor proteins on the surface of the host cell. Importantly, none of the phage genotypes examined that had just three of the four required mutations had any capacity whatsoever to infect hosts that expressed only the OmpF receptor. It would be extremely unlikely to detect a new function that required a quadruple mutation using traditional microbiological methods. However, Meyer *et al.* (2012) observed the mutations accumulate over time as the phage and bacteria coevolved. In fact, the new function emerged repeatedly and quickly, arising in 24 out of 96 replicate populations after only 12 days on average. The question arises, then, as to how the phage evolved this new function, and so quickly, if the intermediate genotypes did not confer any ability to exploit the alternative receptor.

One compelling hint was found in the highly non-random pattern of substitutions observed among the independently derived *J* alleles. In particular, all of the phage that could infect through the OmpF receptor had mutations in four narrow regions of the *J* gene (Meyer et al. 2012). The four mutations did not arise simultaneously because alleles with one, two, or three of these mutations were sampled at earlier time points (Meyer et al. 2012). The genetic parallelism evident in the intermediate states strongly suggested that natural selection favored the intermediate *J* alleles, even though they did not allow the phage to use the OmpF receptor. Meyer *et al.* (2012) also observed that these phage mutations accumulated while the hosts were evolving substantial, but not complete, resistance through reduced expression of the LamB receptor. This synchrony suggested that the reduction in the density of the LamB receptor increased the strength of selection on λ to improve its binding to the ancestral receptor, thereby favoring certain mutations in *J* that, fortuitously, served as necessary stepping stones on the way to the new capacity to use the OmpF receptor. Under this scenario, the arms race between the coevolving hosts and parasites changed the fitness landscape for the phage populations, thereby opening new – and uphill – paths to the innovation (Thompson 2012; Meyer et al. 2012).

Taken together, these observations and hypotheses lead to two testable predictions that are the foci of our paper. First, they predict that the mutations that characterize the intermediate phage types confer a fitness advantage. Second, and more specifically, these genotypes should possess a greater advantage when competing for coevolved hosts than when competing for ancestral hosts; in the extreme case, the intermediate phage types might have an advantage only on the coevolved host cells. To test these predictions, we measured the fitness of six independently evolved phage λ intermediates in competition with the ancestral phage for either sensitive ancestral or partially resistant coevolved host cells.

## Materials and Methods

### OVERVIEW

In the coevolution experiments performed by Meyer *et al.* (2012), a strictly lytic strain of phage λ, called cI26, and *E. coli* B strain REL606 (Table S1) were propagated together in 96 replicate communities by transferring 0.1 ml from each 10-ml culture into a new flask containing 9.9 ml of fresh medium every day for 20 days. Before each transfer, a separate 0.2-ml sample was stored frozen; these samples provided viable representatives of the λ and *E. coli* populations that were used to reconstruct the evolutionary dynamics. Meyer *et al.* found that the first mutations to reach high frequency in the bacterial populations were in *malT*, which codes for a positive regulator of *lamB*, which in turn encodes the native phage λ receptor protein LamB. These mutations reduced expression of LamB, thereby conferring substantial (but not complete) resistance to the ancestral phage, and they reached near-fixation in all host populations by the eighth day of the experiment. Also, one-quarter (24/96) of the phage populations evolved the new ability to infect the bacteria through the OmpF protein, which took 12 days on average (range: 9-18 days).

To study the fitness of the intermediate λ genotypes, we sampled a single phage isolate from each of six independently evolved populations; each isolate was sampled exactly four days before the ability to use OmpF was first observed in that population. The six populations were chosen haphazardly, except that attention was paid to choosing phage from replicates that had evolved distinct *J* alleles by the end of the original experiment. The six genotypes were sequenced to confirm that they had some, but not all, of the four mutations needed to exploit OmpF. Relative fitness was measured in a manner similar to that used in studies on *E. coli* evolution (Wiser et al. 2013), where evolved genotypes compete head-to-head against a genetically marked ancestor. Fitness was assessed in two environments that differed only in the identity of the bacterial host genotype. In one environment, the host was the sensitive ancestral strain, REL606; in the other, the host was an evolved, partially resistant strain, EcC4, that has a single point mutation in *malT* that generates a premature stop codon early in the gene at amino-acid position 295.

### CULTURE CONDITIONS

All *E. coli* cultures were inoculated from freezer stocks into Luria-Bertani (LB) broth (Sambrook and Russell 2001), then grown overnight at 37°C while shaking at 120 rpm. Host cells were preconditioned for the phage competition experiments by growing them in the same medium and other conditions used in the coevolution experiments performed by Meyer *et al.* (2012): 10 ml of modified M9 (M9 salts with 1 g/L magnesium sulfate, 1 g/L glucose, and 0.02% LB) for 24 h at 37°C and 120 rpm. Phage strains were isolated by plating diluted frozen samples on a lawn of REL606 cells suspended in soft agar (LB with 0.8% w/v agar) and spread on the surface of an LB plate (LB broth with 1.6% w/v agar). We then picked an individual plaque – a small clearing in the host lawn caused by the phage population expansion from a single infection – for each phage strain of interest. These isolates were immediately re-suspended in soft agar and the plaque-picking procedure was repeated to ensure that only a single phage genotype was sampled. Next, the phage stocks were grown on ∼10^8^ cells of REL606 for ∼16 h in the same modified M9 medium and other conditions as above. The phage were then harvested by chloroform preparation (Adams 1959) and stored at 4°C. Samples of each isolated phage were also preserved at -80°C in 15% v/v glycerol.

### SEQUENCING THE *J* GENE

We sequenced the *J* gene (host specificity protein, GenBank: NP_040600) from all six λ isolates because previous work showed that the mutations responsible for the new ability to exploit the OmpF receptor were in this gene (Meyer et al. 2012). The sequencing was done using an ABI 3730xl by the Michigan State University Research Technology Support Facility. To that end, PCR-amplified fragments of the *J* gene were sequenced after column-purification with the GE Illustra GFX kit; Table S1 shows the primers used for amplification and sequencing. Point mutations were automatically scored using the DNASTAR SeqMan SNP analysis tool and then manually confirmed.

### FITNESS ASSAYS

We performed fitness assays by setting up direct competitions between evolved λ isolates and a genetically marked ancestor that can be differentiated when grown on a lawn of *E. coli* cells. Some aspects of the assays had to be refined before reliable estimates could be obtained, and they are discussed in the Supporting Information. Here we provide the methods used in the final approach and some background on protocol development.

First, we modified the ancestral λ strain to produce blue, rather than clear, plaques on a lawn of *lacZα*^-^ *E. coli* (strain DH5α) supplemented with X-gal (5-bromo-4-chloro-3- indolyl-β-D-galactopyranoside). When *E. coli* cells metabolize X-gal, they produce a blue compound; however, cells without a functional *lacZα* gene cannot metabolize X-gal. By inserting a functional *lacZα* gene into the phage genome, the phage complements the cell’s capacity to metabolize X-gal and generates a blue spot where a plaque forms. To construct a marked version of phage λ strain cI26, we infected *E. coli* cells that contained a plasmid encoding the phage *R* gene fused to *lacZα* (Wang et al. 2003; Shao and Wang 2008). Phage λ possesses an efficient system for homologous recombination (Ellis et al. 2001), allowing some of the resulting genomes to recombine with the *R*-*lacZα* fusion. The phage progeny were then plated on a lawn of DH5α hosts in the presence of X-gal, and a single blue plaque was chosen. Upon studying this marked strain, we found that the marker was not neutral but instead reduced phage fitness. To quantify the marker’s fitness cost, we computed the mean ± 95% confidence intervals of the selection rate against the marker across several blocks of experiments on each host (ancestral host REL606: 0.120 h^-1^± 0.019 based on six blocks; evolved host EcC4: 0.039 h^-1^ ± 0.017 based on four blocks). The marker effect varied by host (F_1, 54_ = 46.4, *p* < 0.001) but not block (*F*_5, 54_ = 1.395, *p* = 0.241). A fitness cost was also reported for this marker in a related λ strain when phage competed for *E. coli* K12 hosts (Shao and Wang 2008).

Competition assays were run in the same medium and other conditions as used in the coevolution experiments performed by Meyer *et al.* (2012). For each of the six phage isolates tested, we grew one stock culture under the conditions described above. We used aliquots from that stock culture to run replicate competition assays on both the ancestral (REL606) and evolved (EcC4) hosts. Competitions were done in sets of eight assays per host, except for phage B2, which was run with four assays per host; procedural errors led to occasional missing values. The assays were blocked such that all replicates for one phage genotype were assayed on both hosts on the same day. To initiate a competition, we added ∼10^8^ *E. coli* cells (REL606 or EcC4) acclimated to the culture conditions, ∼10^5^ of the marked ancestral phage λ_lacZ_, and ∼10^5^ of a particular evolved λ genotype to each flask. The host density was chosen to match the start of the coevolution experiments performed by Meyer *et al.* (2012); however, phage densities were almost 10-fold lower because we could not produce dense stocks of the evolved genotypes, despite several attempts. (The lower density should reduce competition between the phage genotypes and thereby diminish our ability to detect fitness differences. In fact, however, we measured fitness gains for the evolved phage, as shown in the Results.) The mixed cultures were incubated for 8 h, and the phage were sampled and enumerated at the beginning and end of that period. We chose a duration of 8 h for two reasons: (i) given the large fitness advantages of the evolved genotypes, they would drive the ancestral phage to very low density if the assays ran for 24 h, thereby compromising our ability to quantify the fitness difference; and (ii) resistant host mutants often arose and reached substantial frequency in 24 h (Supporting Information text), thus altering the competitive environment of the phage. Phage samples were diluted in TM (50 mM tris hydrochloride, pH 7.5 and 8 mM magnesium sulfate), and multiple dilutions were plated to find one where individual plaques could be counted. Plates used to count the phage contained DH5α host cells and 9.5 mg ml^-1^ X-gal in the top agar, and they were incubated at 37°C for two days to allow the blue plaques to develop full pigmentation.

We calculated fitness as the difference, not ratio, of the two competitors’ realized growth rates during the competition. This quantity, sometimes called the selection rate *r*, is related to the more familiar ratio of the growth rates (i.e., relative fitness *W*). However, we used the difference here because it is more robust to large differences in growth rates between competitors, including those cases where one competitor declines and the other increases in abundance, (Lenski et al. 1991; Travisano et al. 1995), which we sometimes observed.

### STATISTICAL ANALYSES

We expected the fitness of the evolved phage (expressed as a selection rate differential relative to the marked ancestral phage) to be greater on the coevolved EcC4 host cells than on the ancestral cells. We also expected the realized growth rates of the evolved phage to be lower on the coevolved than on the ancestral cells. Therefore, we employed one-tailed *t-*tests of these hypotheses. (In those cases where the outcome was opposite to our expectation, we report the p-value as >0.5.) We tested each hypothesis for six λ genotypes, and therefore we performed Bonferroni corrections for multiple comparisons, resulting in an adjusted α = 0.05/6 = 0.0083 per test. We performed a single *t*-test with α = 0.05 to assess the effect of the coevolved versus ancestral phage competitors on the realized growth rate of the ancestral phage.

We performed a two-way ANOVA to determine the effect of assay date (block) and host type (coevolved versus ancestral) on the fitness cost of the phage’s *lacZα* marker. This analysis showed that the *lacZα* marker had a more deleterious effect on phage fitness when competing for the ancestral cells than for the coevolved *malT*¯ cells. Therefore, we tested whether correcting for this effect would affect our interpretation of the differences in phage fitness on the two host types; the correction was a simple subtraction of the grand mean of the difference in the fitness cost of the marker between the two hosts. In fact, this correction did not affect our interpretation (Table S2).

All analyses were conducted in R (R Development Core Team 2011) using custom scripts.

## Results and Discussion

### MUTATIONS IN THE *J* GENE OF THE SIX PHAGE ISOLATES

The six independently derived λ genotypes that we studied had different subsets of the four *J* mutations needed to exploit the OmpF receptor (Table 1). In addition, all six had one or more other mutations in the *J* gene that were not required to use OmpF (Table 1). In four cases (A7, B2, D9, and G9), the additional mutations were also present in the eventual OmpF^+^ phage, indicating that these genotypes were true intermediates on the evolutionary paths leading to the OmpF^+^ phage. In two cases (A12 and E4), the other mutations were not present in the later OmpF^+^ isolates, implying that these genotypes were somewhat off the direct line of descent leading to the OmpF^+^ phage.

**Table 1.** Mutations in the *J* gene of the six evolved phage λ isolates in this study. Each virus was randomly sampled from a different source population four days before phage that could use the OmpF receptor were first detected. Each mutation is identified by the ancestral nucleobase, its position in the reading frame of the *J* gene, and the evolved nucleobase. Asterisks under “Line of Descent” indicate that the corresponding mutation was present in the later phage able to use the OmpF receptor. Previous research (Meyer et al. 2012) showed that phage λ requires four mutations—one or more in four specific regions of the *J* gene—to exploit the OmpF receptor. The mutations in bold are located in these regions; the last column shows how many of the four requisite regions harbored mutations. All values are less than four because these isolates were intermediates not able to use the OmpF receptor. Isolates A7 and G9 have three bolded mutations each, but only two of the mutational categories are satisfied because the first two bolded mutations affect the same region of the *J* gene.

| Source Population | Day of Isolation | Set of Mutations | Line of Descent | Number of OmpF Categories Satisfied |
| --- | --- | --- | --- | --- |
| A12 | 10 | C599T | * | 1 |
|  |  | G2921A |  |  |
|  |  | **T2991G** | * |  |
|  |  | C3033T |  |  |
|  |  | C3147A |  |  |
|  |  | T3380C | * |  |
| A7 | 10 | **A2989G** | * | 2 |
|  |  | **C2999T** | * |  |
|  |  | C3119T | * |  |
|  |  | **G3319A** | * |  |
| B2 | 13 | **C2969T** | * | 2 |
|  |  | C3119T | * |  |
|  |  | **G3319A** | * |  |
| D9 | 8 | C2879T | * | 2 |
|  |  | **T2991G** | * |  |
|  |  | C3119T | * |  |
|  |  | **G3319A** | * |  |
| E4 | 13 | **C2969T** |  | 2 |
|  |  | C3119T | * |  |
|  |  | **G3319A** | * |  |
| G9 | 11 | A1747G | * | 2 |
|  |  | **A2989G** | * |  |
|  |  | **C2999T** | * |  |
|  |  | C3119T | * |  |
|  |  | **G3319A** | * |  |

### MUTATIONS IMPROVE FITNESS PRIOR TO ORIGIN OF NEW FUNCTION

All six evolved phage isolates were more fit than their ancestor when competing for the coevolved host cells, and four of them (A12, A7, D9, and G9) were more fit than their ancestor on both the ancestral and coevolved host cell types (Figure 1). These data are the first direct experimental evidence that these intermediate phage genotypes had selective advantages even before they had evolved the ability to exploit the OmpF receptor. Such benefits were previously hypothesized because, without them, it was unclear how the several mutations required for phage λ to evolve the ability to exploit the OmpF receptor had accumulated so quickly and repeatedly across the replicate populations (Meyer et al. 2012). These data show that the intermediate steps leading to the new function were on balance favored by selection, and thus they help to explain how the phage achieved that innovation.

**Figure 1.**
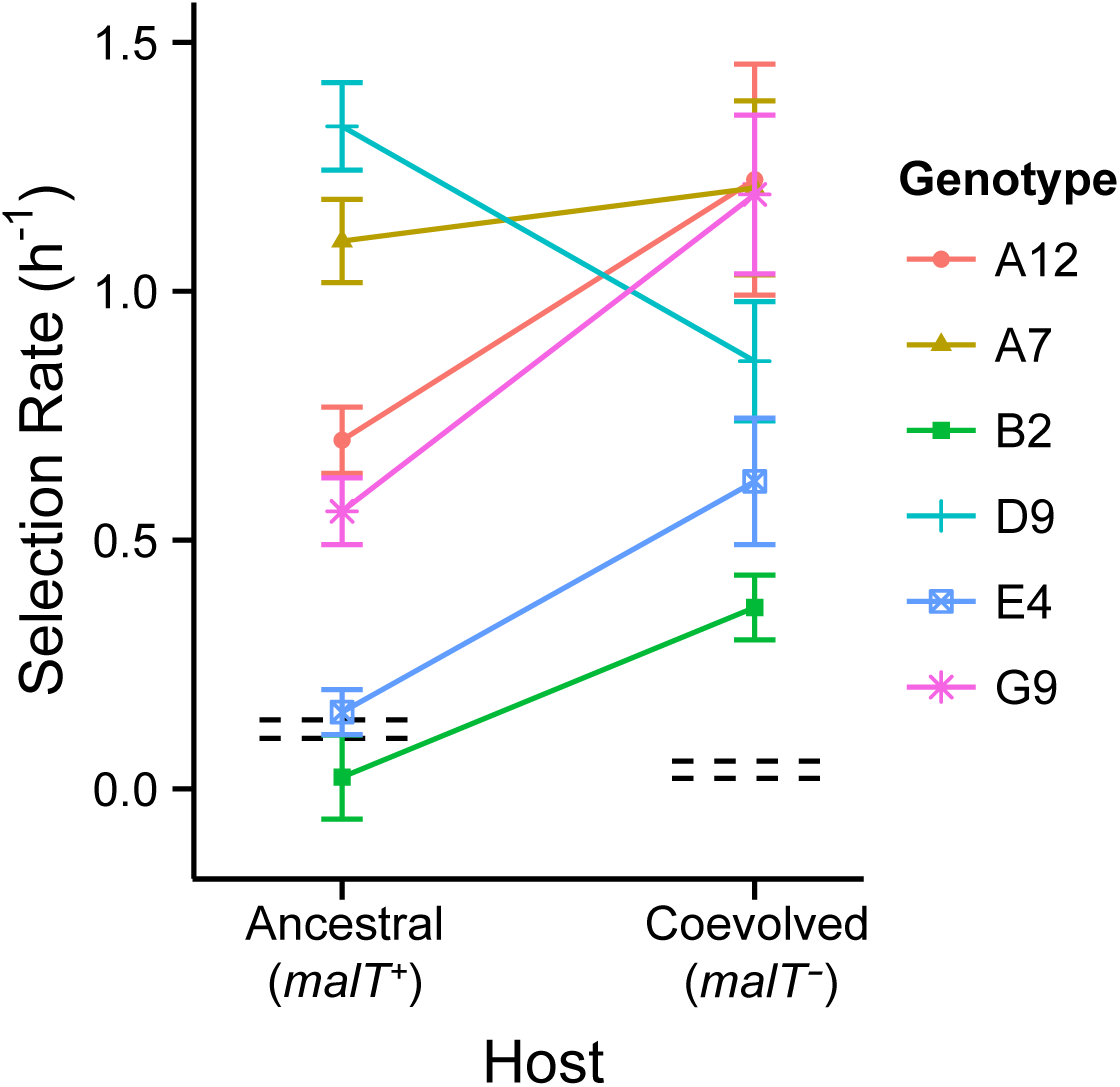
Relative fitness of evolved phage λ genotypes on ancestral *malT*^+^ (REL606) and coevolved *malT¯* (EcC4) hosts. All fitness values were measured in competition with a genetically marked variant of the ancestral phage. Error bars around points are 95% confidence intervals. The marker imposed a small fitness cost on the phage that differed for the two hosts; the black dashed lines indicate the 95% confidence intervals for the fitness of the unmarked ancestor relative to the marked variant. Four of the six evolved phage genotypes (A12, A7, D9, and G9) were significantly more fit than their ancestor on the ancestral host; all six were significantly more fit than their ancestor on the coevolved host. Also, four of the evolved phage genotypes (A12, B2, E4, and G9) had significantly higher relative fitness on the coevolved host than on the ancestral host (Table S2).

### HOST COEVOLUTION AFFECTS VIRUS FITNESS

In five of the six cases analyzed here, the fitness of the evolved phage genotypes relative to the ancestral phage differed depending on whether they competed for the ancestral *malT*^+^ host or the coevolved *malT*¯ host (Figure 1). In four cases – A12, B2, E4, and G9 – the advantages of the evolved phages were significantly higher during competition for the coevolved hosts. (The advantage of evolved phage A7 was also slightly higher on the coevolved host, although that difference was not significant.) By contrast, evolved phage D9 had a greater advantage when competing for the ancestral host than when competing for the coevolved host. Thus, in four of the cases examined here, coevolution of the host population accelerated the evolution of phage genotypes on the path leading to the origin of the new ability to use the OmpF receptor.

We can gain further insight into the evolutionary changes in the phage by looking at the underlying growth rates of the evolved and ancestral phage populations during the competition assays (Figure S1). Five of the six evolved phages – all except G9 – grew more slowly during the competitions on the evolved *malT¯* host than they did during the competitions on the ancestral *malT*^+^ host. However, the ancestral phage could not grow to any measurable extent on the evolved *malT¯* host (Figure S1). As a consequence, all six evolved phage genotypes were more fit than their ancestor in competition for that host type (Figure 1), even though the evolved phage grew more slowly on that host type. By contrast, the ancestral phage grew well on the ancestral *malT*^+^ host (Figure S1), so that only four of the evolved phage genotypes (A12, A7, D9, and G9) were significantly more fit than their ancestor in competition for the ancestral host (Figure 1).

Taken together, our results reveal important similarities and substantial variation among the six intermediate phage types examined here. Most importantly, and consistent with the general hypothesis that the intermediates reached high frequencies because the mutations they carried conferred selective advantages, all six phage were significantly more fit than their ancestor when competing for the ancestral host, the coevolved host, or both. The results also tend to support, though not in every case, the specific hypothesis that host coevolution pushed the evolving phage populations along paths that accelerated the rise of the intermediate types and hence the subsequent emergence of phage with the novel ability to exploit the OmpF receptor. This latter trend is consistent with theory that predicts coevolution can drive innovation by altering the shape of fitness landscapes in ways that favor new phenotypes (Rosenzweig et al. 1987; Weitz et al. 2005; Williams 2013). Several other experimental studies also support the hypothesis that coevolution can promote evolutionary innovation. Populations of phage *ϕ*2 explored genomic space more broadly when they adapted to coevolving hosts than when they adapted to a static host (Paterson et al. 2010). When phage M1, a relative of T2, was sequentially presented with hosts expressing potential receptors that were increasingly different from its usual receptor, the phage evolved the ability to use a protein not previously known to have that capacity (Hashemolhosseini et al. 1994). Also, recent experiments with digital hosts and parasites, analogous in some respects to bacteria and phage, showed that coevolution generated negative frequency-dependent selection that deformed fitness landscapes in ways that often favored the evolution of new, more complex traits (Zaman et al. 2014).

### COMPETITION FOR RECEPTORS DRIVES EVOLUTION OF J PROTEIN

Phage λ uses its J protein to adsorb to the LamB receptors on the surface of host cells (Gurnev et al. 2006), and the mutations that distinguish the intermediate phage genotypes from their ancestor are in the *J* gene that encodes that protein. Therefore, it seems likely that differences in the ability to recognize and adsorb to the LamB receptors cause the observed fitness differences among the λ genotypes. In particular, this hypothesis would explain why the intermediate types had a fitness advantage even before they acquired the new ability to exploit the alternative OmpF receptor. This hypothesis might also explain why most of the intermediates have larger advantages when competing for the coevolved *malT¯* hosts, which differ from the ancestral host by expressing fewer LamB receptors.

To test for the effect of competition for the host-cell receptors, we compared the realized population growth rate of the marked ancestral phage during competition against either the unmarked ancestor or the evolved phage intermediates with mutations in their J proteins. If competition for LamB receptors drove J evolution, then we would expect the evolved phage to be better at finding the receptors and thereby occluding the ancestral phage from binding to them. As a consequence, the evolved phage would also initiate infections more quickly and reduce the number of host cells available for subsequent rounds of infection. Indeed, the marked ancestral phage grew significantly slower in the presence of evolved phage competitors than in the presence of other ancestral phage on the ancestral *malT*^+^ host (*t* = 1.891, one-tailed *p* = 0.044, Figure 2). We could not perform a similar analysis on the coevolved *malT*¯ host, however, because the ancestral phage did not grow at all during those competition experiments (Figure S1).

**Figure 2.**
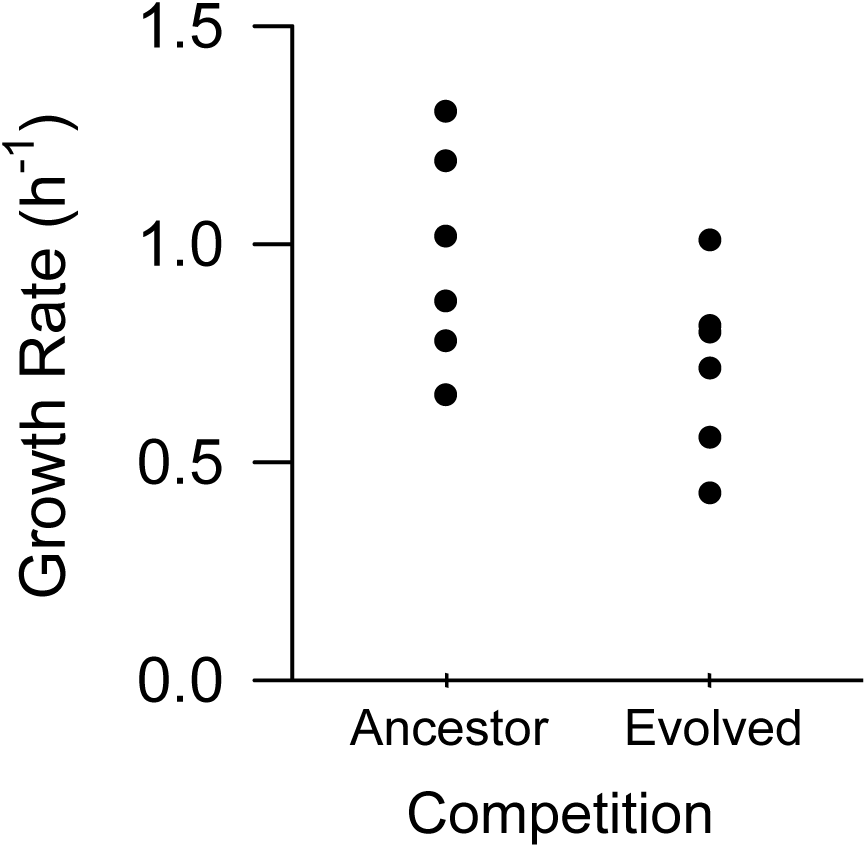
Realized growth rates of the ancestral phage depend on the competition. The marked ancestral phage grew more slowly on the ancestral host in the presence of evolved phage competitors than in the presence of its unmarked ancestral counterpart. Each point shows the mean of one set of assays; each set was performed on a different date, and the number of competitions in each set varied from four to eight. The realized growth rates measured against the evolved phage competitors present the same data as shown in Figure S1 (for the ancestral host only).

Competitive displacement can be an important driver of parasite evolution. For example, a pandemic strain of influenza was observed to displace two seasonal influenza strains during coinfection of host cells (Perez et al. 2009). During competition for aphid hosts, the parasitic wasp *Aphidius ervi* competitively displaced its competitor *A. smithi*, even though *A. smithi* searched more efficiently for hosts when it was alone (Chua et al. 1990). In a direct test of the importance of competitive displacement on phage evolution, *ϕ*6 was shown to evolve the ability to infect different *Pseudomonas* species sooner when the ratio of viral particles to host cells was increased, thereby intensifying competition (Bono et al. 2012). Together these studies demonstrate the importance of competition for the evolution of a parasite’s ability to exploit hosts. Moreover, they show how changes in the competitive environment – whether from increased parasite density (Bono et al. 2012) or, as we have shown here, improved fitness of competing parasites – can influence parasite evolution. Therefore, the adaptive landscape of a virus or other parasite can be reshaped not only by changes in the genetic composition and density of its hosts but also by changes in the density and fitness of its competitors for those hosts.

### IMPLICATIONS OF ECOLOGICAL INTERACTIONS FOR PREDICTABILITY OF EVOLUTION

It is important to account for the effects of ecological interactions, including with both conspecifics and other species, on the relative fitness of different genotypes if one is to understand and even predict evolutionary changes (Bohannan and Lenski 2000; Meyer and Kassen 2007). Here we have quantified how a host’s coevolution led to varied – and sometimes opposite – effects on the relative fitness of virus genotypes (Figure 1). We also showed that the growth rate of a given virus population depended on the genotypes of its competitors (Figure 2). Predicting the course of viral evolution therefore depends on knowing the genetic states of both the host and parasite populations and understanding how those states affect fitness. Scaling this understanding to more diverse and complex ecosystems will be challenging (Telfer et al. 2010), but it may be necessary to predict such phenomena as disease emergence. The fact that coevolution generates dynamic fitness landscapes also implies that populations are more evolvable than they would otherwise be. As a consequence, key innovations and other hard-to-reach adaptations may be more likely to evolve in these dynamic landscapes.

## Acknowledgments

We thank Neerja Hajela and Rachel Sullivan for assistance in the laboratory, and Ben Kerr, Jenna Gallie, and Kayla Miller for comments on earlier versions of this paper. This research was supported by the BEACON Center for the Study of Evolution in Action (National Science Foundation Cooperative Agreement DBI-0939454), a National Science Foundation Graduate Research Fellowship to ARB, and the John Hannah endowment from Michigan State University.

